# Positive associations matter: microbial relationships drive tick microbiome composition

**DOI:** 10.1101/2022.11.06.515366

**Authors:** Nicholas M. Fountain-Jones, Benedict S. Khoo, Austin Rau, Jesse D. Berman, Erin N. Burton, Jonathan D. Oliver

## Abstract

Untangling how factors such as environment, host, associations between species and dispersal predict microbial dynamics is a fundamental challenge. In this study, we use a robust sampling design coupled with complementary machine-learning approaches to quantify the relative role of these factors in shaping microbiome variation of the blacklegged tick *Ixodes scapularis. I. scapularis* is the most important vector for *Borrelia burgdorferi*. (the causative agent for Lyme disease) in the U.S as well as a range of other important zoonotic pathogens. Yet the relative role of the interactions between pathogens and symbionts compared to other ecological forces is unknown. We found that positive associations between microbes where the occurrence of one microbe increases the probability of observing another, including between both pathogens and symbionts, was by far the most important factor shaping the tick microbiome. Microclimate and host factors played an important role for a subset of the tick microbiome including *Borrelia* (*Borreliella*) and *Ralstonia*, but for the majority of microbes, environmental and host variables were poor predictors at a regional scale. This study provides new hypotheses on how pathogens and symbionts might interact within tick species, as well as valuable predictions for how some taxa may respond to changing climate.

## Introduction

The intricate ecological drivers of within-host microbial communities are poorly understood for most species. For many species, environmental factors such as temperature are well-known drivers of host behaviour and distribution, but these factors can also directly or indirectly shape the composition of the microbial communities that inhabit them [e.g., 1–5]. Host factors such as sex and age as well as the spatial relationships between hosts are increasingly considered important in determining within-host microbial communities (hereafter ‘microbiomes’) [e.g., 6–8]. For example, hosts closer in space tend to have more similar microbiomes compared to those further away indicating that microbial dispersal can be important in structuring animal microbiomes [e.g., 7]. Hormonal and development differences between sexes can also lead to changes in microbiome composition [6, 9]. However, the impact of associations between microbes is less often quantified but can explain substantial amounts of microbiome variation [8, 10, 11]. For example, environmental effects on *Ixodes ricinus* microbiome dynamics were relatively weak yet significant positive and negative associations between microbes were detected (i.e., species that co-occur more or less than expected by chance respectively) [10]. Positive associations can be proxies for apparent facilitation (i.e., colonization by one species facilitates another) and negative associations can be proxies for competition between taxa [12] (however see [13]). Non-random associations between bacteria in the human gut have been found to be often negative which likely indicates that competition is an important driver of structure in this community [14]. However, the relative importance of environment, host and microbial associations in microbiome organization and assembly is unknown for most species, including arthropod vectors of pathogens.

The microbiome of vector species is increasingly seen as playing an important role in shaping vector-borne pathogen dynamics in arthropod vectors, such as ticks (Arthropoda: Ixodida). Ticks are one of the most important vectors of pathogens affecting humans and carry a wide range of pathogenic bacteria such as *Borrelia burgdorferi* (the causative agent for Lyme disease) and *Anaplasma phagocytophilum* (the causative agent for human granulocytic anaplasmosis). The non-pathogenic members of the ticks microbiome can have variable effects on pathogens and this can, in turn, have impact human health [15, 16]. For example, in the deer tick *I. scapularis*, maternally inherited *Rickettsia* species may limit the colonization of *B. burgdorferi* and cause lower infection rates [15]. Artificial introduction of the symbiotic bacterium *Wolbachia* to *Anopheles* mosquito has been shown to interfere with *Plasmodium* species (the causative agents for malaria) and limit infection and onward transmission to humans [e.g., 17]. Symbionts can also promote pathogen infection as, for example, ticks reared without external symbiont colonization took larger blood meals and had reduced *B. burgdorferi* loads compared to controls [18]. However, tick microbiomes are complex and for *I. scapularis* consist of hundreds of potentially interacting taxa just in the U.S [e.g., 19, 20]. Experimental approaches to untangle these interactions is impractical and can include interactions not found in the wild [16].

Quantifying associations in the wild and their relative importance compared to other ecological drivers is a methodological challenge [10, 21]. For example, it is important for controlling for spatial and temporal patterns as species may appear to cooccur less frequently than expected by chance because they are spatially segregated or limited by dispersal rather than through microbial competition [22–24]. Positive associations may be a product of shared habitat preferences so careful consideration is needed to include environmental factors appropriately [22]. Graphical network models such as conditional random fields (CRFs) can start to untangle the complex dynamics shaping microbiome assembly by estimating microbial associations between microbiome constituents while accounting for spatial, host and environmental effects [8, 25]. CRF models also capture complicated random effect structures that can quantify site effects that are common in tick microbial communities. Compared to joint species distribution models (JSDMs) such as HMSC [26], CRF directly estimates the conditional effect of one species’ presence on another’s occurrence probability (while accounting for spatial and environmental effects) [25]. One weakness of this approach, however, is that CRF models do not capture non-linear relationships between environmental variables and microbial communities or provide a framework to interrogate these relationships further. The recently developed multi-response interpretable machine learning (MrIML) R package [27] offers exciting tools that can capture and interrogate nonlinear host-environment relationships within microbiomes. While currently unable to assess microbial associations or model the effect of distance between sites, MrIML models provide insight into what role host and environment plays in shaping microbiome composition and identify individual taxa most strongly responding to a particular gradient. Similar modelling approaches have uncovered some of the complex environmental relationships shaping plant and animal communities [27, 28] and genetic turnover [29] but are rarely used to quantify the non-linear drivers of microbiome composition.

In this paper we couple a robust bioinformatic pipeline and innovative analytical framework leveraging both MrIML and graphical network models to untangle how microbial associations, environment, and host shape the microbiomes of *I. scapularis* ticks across a climate gradient in the U.S. Midwest (Fig. 1). While quantifying the drivers of microbiome organization, we also examine the drivers of focal members of the community, including potential pathogens (Table 1). We demonstrate that approaches such as ours can begin to decipher microbiome dynamics and also guide experimental efforts to test microbial associations that can be crucial in disease control efforts [30].

**Fig 1.**
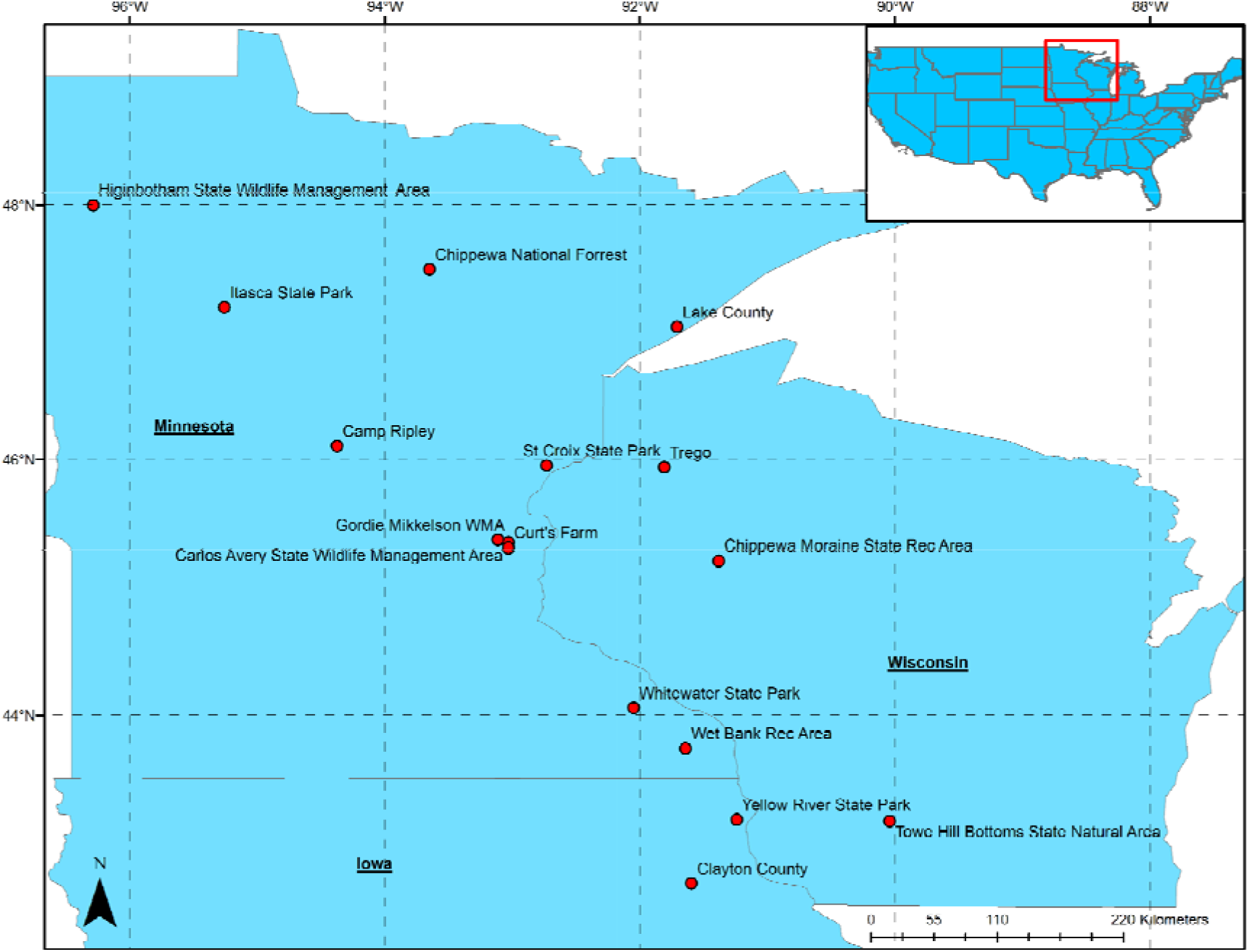
Map of tick sampling sites across Minnesota, Wisconsin and Iowa. Red box in the upper smaller map shows geographical location in relation to the 48 lower continental U.S. states.

**Table 1:**
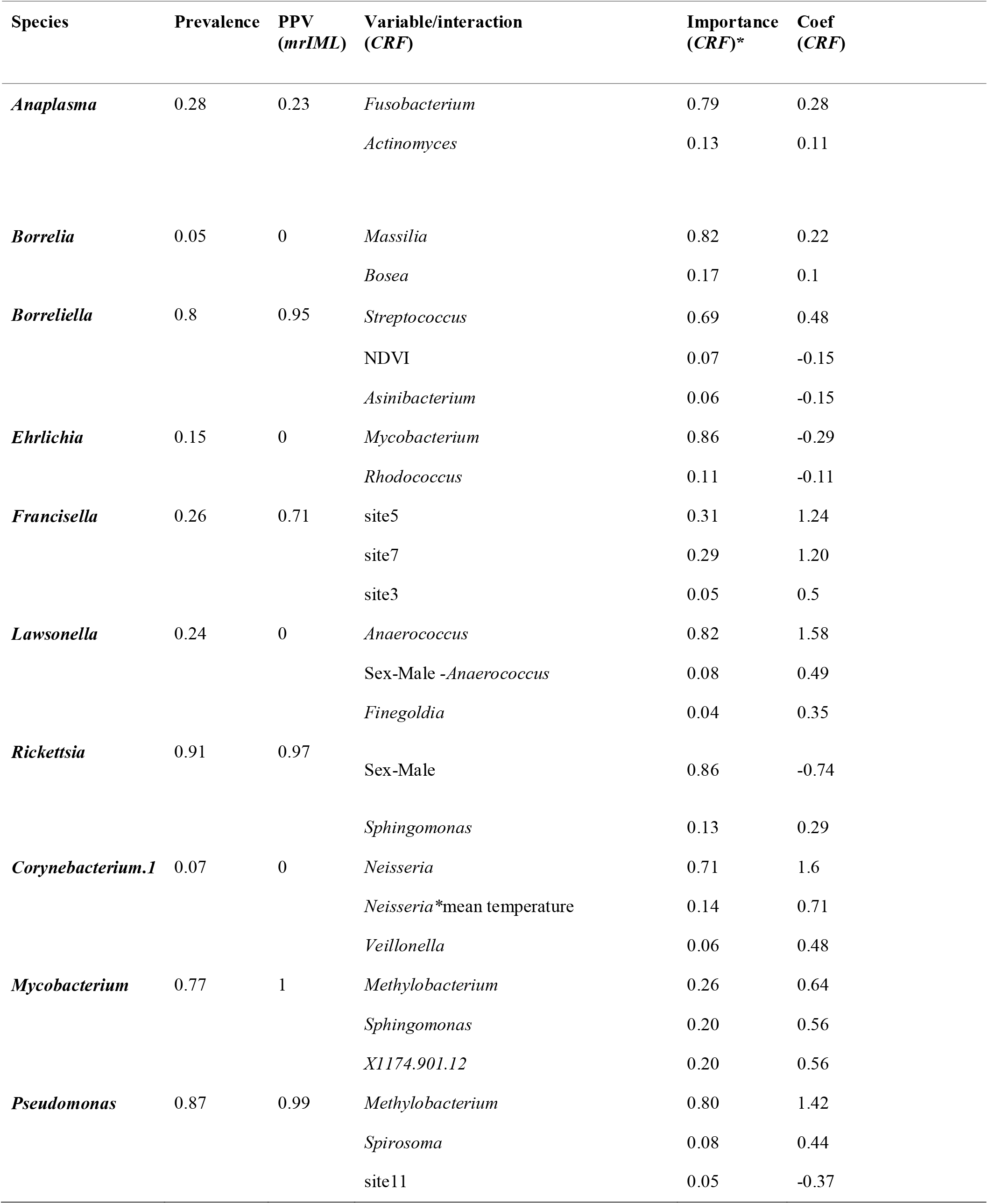
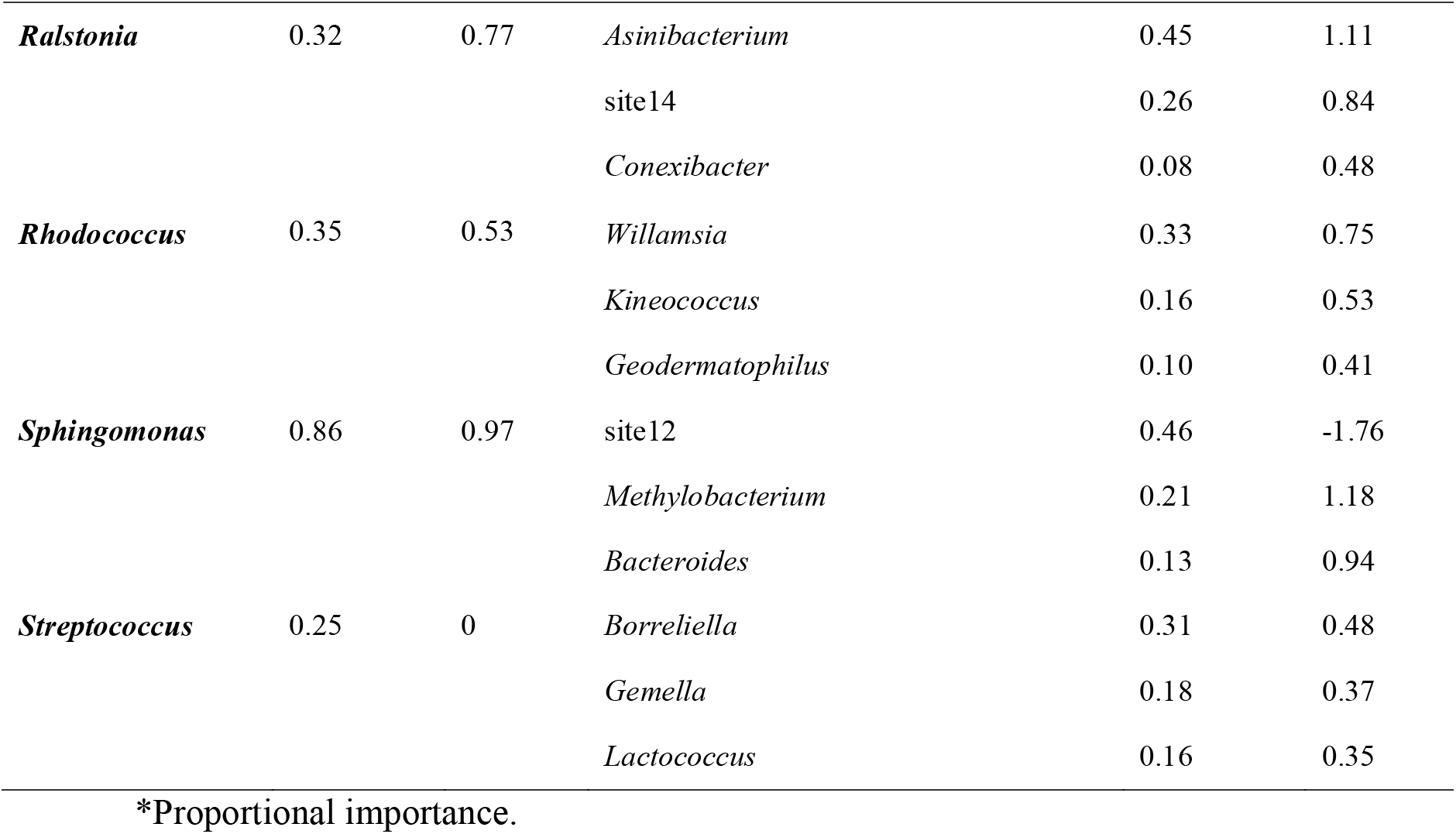
Model variable importance and standardised coefficient estimates for the top three variables with coefficient estimate > ±0.1 for our focal taxa.

## Methods

### Sample collection

Tick samples were collected using the dragging method from 16 sites in the US Upper Midwest between 2017-2019. Collection sites were selected along a North/South gradient (Fig. 1). Only adult ticks were included in this analysis. While still living, all ticks were washed to reduce surface contaminating bacteria as much as possible [31]. Briefly, ticks were immersed in 10 ml distilled water + 1 drop of polysorbate 20 (agitated for 5 min), immersed in 10 ml 0.5% benzalkonium chloride (agitated 5 min), immersed in 10 ml 70% ethanol (agitated 5 min), rinsed in 10 ml water (agitated 1 min), and placed in 70% ethanol for storage at room temperature with blotting to remove moisture between all steps. Ticks were bisected and incubated in a 180 μl lysis buffer at 60 °C overnight. While incubating, samples were vortexed multiple times [32]. DNA extraction was performed using DNeasy Blood and Tissue kits (Qiagen, Hilden, Germany) according to the manufacturer’s instructions.

Soil samples were collected from three locations at each site to a depth of 15 cm using a soil probe, with samples taken at least 5 metres apart from each other. Soil samples were homogenised, and DNA was extracted using DNeasy PowerSoil Pro kits (Qiagen) according to the manufacturer’s instructions.

Appropriate permits and permissions were collected from all collection sites where required.

### 16S microbiome analysis and taxonomic designation

DNA samples were then plated on 96-well plates with 2 wells on each plate being left for a blank and a negative control. Additionally, two out of the five total 96-well plates had spatter control and plate effect controls added in. Spatter control samples consisting of 5 wells with extracted *Brachyspira pilosicoli* DNA that were strategically spread through the plate, with one of the two plates having 2 additional blank wells spaced near these spatter controls act as spatter standards. *B. pilosicoli* DNA was used because it was judged to be very unlikely to occur at the tick collection sites as the upper Midwest US lacks a wild pig population and *B. pilosicoli* as an enteric pathogen cannot be up taken by ticks during blood meals. The plate effect controls consisted of 3 samples from the other three plates each. Sequencing of the V4 region of the 16S was done with an Illumina MiSeq set at 600 cycles.

### Bioinformatics

The resulting 16S reads were then cleaned with cutadapt (https://cutadapt.readthedocs.io) with a minimum length of 100 bases post-removal of sequence adaptors and primers. The DADA2 software package in R was then used for removal, filtering, and trimming of low quality sequences and chimeric reads [33]. The remaining sequence data was then used for error inference and ASV table creation, taxonomic identification of ASVs was performed using the assigntaxonomy command utilising the RDP 18 trainset database. This pathway differentiated *Borreliella* (*Borrelia burgdorferi sensu lato*) from *Borrelia* (relapsing fever borreliae). For the purposes of clarity, we have retained these designations throughout the manuscript. The soil sample microbiomes were evaluated to determine the constituents of the tick microbiome that were likely environmentally acquired, and these were removed from analysis.

Using the OTUtable R package [34], we further filtered out ASVs with less than 1% of reads in a least one sample and occurred in at least in 2.5% of all samples. We chose these cut-off values to reduce class imbalance issues but maximise the numbers of ASVs included in the analysis [8, 27]. We converted the remaining ASV count data into presence/absence data (0/1s). As *Rickettsia* was very common throughout our samples and there was some evidence of splatter across wells, we set threshold with > 600 reads indicating a positive sample and < 600 was negative.

### Environmental data

Daily weather data was acquired from PRISM (Parameter-elevation Relationships on Independent Slopes Model), a blended continuous spatiotemporal climate product [https://www.prism.oregonstate.edu/]. Data was available as 4 km grids that were spatially and temporally overlaid with the 16 tick collection sites. Each collection site was assigned the weather condition of their respective grid cell. Metrics included mean dewpoint, maximum/minimum/mean temperature, and maximum/minimum vapor pressure deficit for the day that ticks were sampled. The average weather conditions were calculated for 30 days, 1-year, 2-years, and 5-years prior to a sampling event for all weather variables. Snowfall data was acquired from the National Snow and Ice Data Center (accessed on 01/12/2020), which provides NASA derived daily snow water equivalents and snow depths at 4 km grids, which were spatially overlaid with collection sites. Two metrics were calculated of 1) maximum snow depth in the winter prior to sampling and 2) number of days in the winter prior to sampling with a snow depth greater than 0 mm [35].

Greenspace was evaluated from the satellite derived time integrated Normalized Difference Vegetation Index (NDVI) using Aqua eMODIS data. Annual NDVI values represent the total photosynthetic activity in a 250m grid during a growing season with higher values indicating a greener more diverse environment. Each collection site was spatially overlaid with the previous year NDVI grid, assigning it a greenspace value from the prior year. Both the NDVI value at the precise collection site location and the average NDVI were extracted within a 2.5 km buffer surrounding each collection site.

### Multivariate interpretable machine learning

Using the environmental and tick sex data, we constructed MrIML predictive models to capture microbiome-wide relationships and identify individual ASVs responding to these predictors. Similarly to the *Gradient Forests* approach [28], MrIML constructs and tunes individual machine learners for each response variable (ASVs in this case) and then summarizes and interprets the combined (‘global’) model [27]. In this way, models constructed with MrIML can capture even strongly non-linear responses to the environment at a community level as well as highlight the taxa most strongly responding to the gradient. The MrIML approach also provides extensive autotuning capabilities to optimize algorithm performance as well as 10-fold cross validation to guard against model overfitting. Moreover, multi-algorithm performance in prediction across ASVs can be easily compared and the model with highest performance further interrogated [27, 36]. We compared the predictive performance of three different algorithms; generalized linear models (GLMs, logistic regression), random forests (RF) and extreme gradient boosting (XGB). For the tree-based methods (RF and XGB), we generated 1000 trees for each ASV and tuned all other hyperparameters using a 5*5 grid. We hot encoded tick sex for the GLM and XGB models. We compared the average positive predictive value (PPV), specificity and sensitivity and also checked variable importance scores (using the “vip” method [37]) for consistency across algorithms. To capture uncertainty in our variable importance measures, we ran the pipeline for 10 iterations with different seeds and summarized variable importance scores. For the best performing model, we computed accumulated local effect plots (ALEs) to assess the conditional effect of the predictor on microbiome composition. ALEs capture the average probability of observing an ASV across predictor values while holding all other variables in the model constant [27 for more details, see 38]. We further identified the taxa responding most strongly to each feature using an ALE standard deviation threshold of 0.1 (i.e., the probability of observing this ASV had to change by a minimum of 10% across the gradient). To examine how the most important variables shaped the occurrence probabilities of *Borreliella* and screen for interactions between variables, we used individual conditional expectation (ICE) plots. Briefly, ICE plots calculate the conditional change in occurrence probability of microbes within each tick (data instance) separately across a grid of variable values constant [27, 38]. ICE plots can, for example, show when the response to environmental or host variables just differs between ticks and an interaction with another variable may be important.

### Graphical networks

To test if including microbial associations as well as space could improve predictions, and account for site and geographical effects we used the graphical network approach outlined in [8, 25]. Our graphical network analysis also allowed us to identify potentially interacting ASVs and assess the relative role of positive and negative interactions in shaping tick microbiome organization across our gradient. To simplify the model, we did not include year in the model as preliminary MrIML models showed that year was of low predictive performance. We compared four graphical models of increasing complexity. The first model separated the effects of only microbial associations in the log-odds of observing each ASV (Markov Random Fields model, MRF), the second model included spatial regression splines to account for spatial autocorrelation between ticks (spatial MRF model), the third included microbial associations, environment, and host effects without spatial splines (CRF model) and the last included all components including spatial splines (spatial CRF model). Briefly, for this class of graphical model, the log odds of observing an ASV are calculated using logistic regression with covariates (CRF) or without covariates (MRF). Cross multiplication of all co-occurring ASV and covariate combinations allow the ecological processes that shape ASV occurrence to be assessed [8, 25]. Edges between ASVs in the co-occurrence network represent conditional coefficient estimates between that pair. We used least absolute shrinkage and selection operator (LASSO) regularization to force coefficients to 0 if they have minimal effects and 10-fold cross validation to choose the penalty that minimized error. This process was repeated 50 000 times for each regression to ensure adequate model fitting. From this data, we constructed an adjacency matrix and plotted association networks using *Igraph* R package using a non-metric multidimensional scaling (nMDS) layout (ASVs closer together share similar edges) [39]. We calculated the degree distribution using *Igraph* with a coefficient threshold ≥ ±0.1. To aid interpretability, we constructed networks with coefficients between taxa ≥ −0.1 for negative associations and ≥ −0.4 for the positive associations. Taxa without associations above these thresholds were removed from the network. See [8] and [25] for more details. See https://github.com/nfj1380/tick_microbiome_analysis for the complete model pipeline.

## Results

### Microbiome diversity and composition

Across our 16 sites, we analysed the microbial communities from 355 ticks We After filtering, we analysed 232 ASVs. We did not see any cross-contamination from sample spatter controls across those 232 ASVs in the wells with tick and soil samples. Nor did we see any significant plate effect that was beyond the error range from the Illumina MiSeq sequencer with our duplicated samples across the 5 plates. The major taxa represented across samples included Rickettsiaceae (purple, Fig. S1), Francisellaceae (green, Fig. S1), and Anaplasmataceae (light brown, Fig. S1).

### Drivers of microbiome variation

Overall, our MrIML models revealed strong heterogeneity in the response of the tick microbiome to environment and host variables. Generally, non-linear relationships better predicted tick microbe responses as the RF-based MrIML model overall had higher predictive performance compared to the GLM-based model (Table S1). XGB had the lowest performance of the three algorithms tested (Table S1). Even though the average specificity and sensitivity of all algorithms was relatively high (0.8-0.95, Table S1), the best-performing RF model could only accurately predict the presence of at ASV on average ~15% of the time (positive predicted value (PPV)= 0.15, Table S1). Our MrIML models could not adequately explain the distribution of 150 out of 232 taxa (PPV > 0.5). However, environmental and host effects could explain the distribution of some microbes remarkably well with PPV >0.7 for 16 ASVs including focal taxa such as *Borreliella* and *Ralstonia* (Fig. 2).

**Fig. 2:**
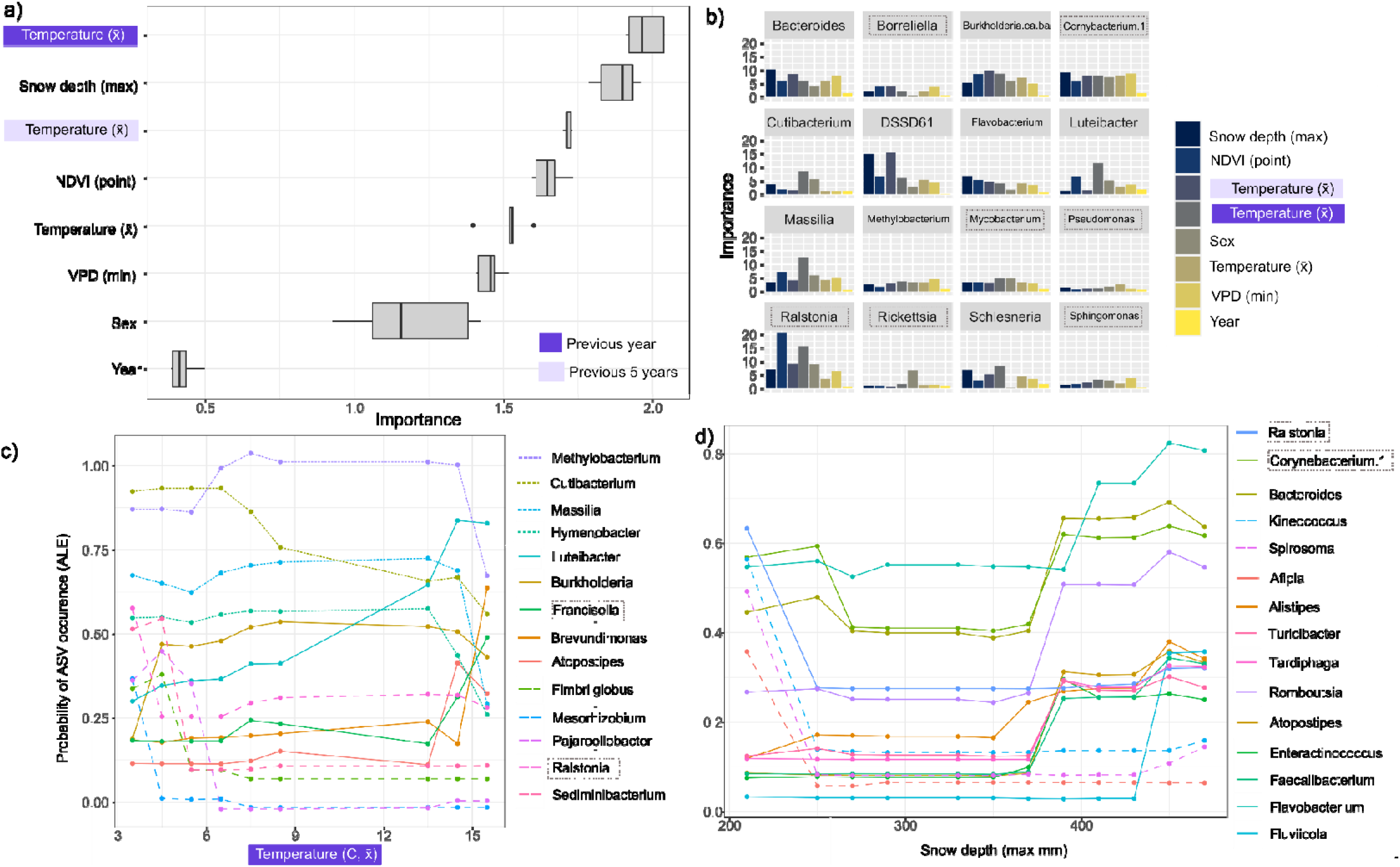
Plots showing the a) microbiome-wide variable importance across multiple runs, b) variable importance for the ASVs best predicted by the model (PPV > 0.7) and c & d) accumulated local effect (ALE) plots showing the conditional relationships for the top two most important predictors for our best performing MrIML microbiome model. Focal taxa are indicated by the dashed boxes. Medium dashed lines =ASVs with a generally negative relationship with the response. Small dashes = decline followed by increase. Unbroken = positive relationship with the response. See Table S1 for model performance estimates across algorithms and Fig S3 for global ALE plots and Fig. S4 for NDVI relationships.

Variable importance was qualitatively similar across MrIML models (Fig. S2) with the previous year’s average temperature the best performing predictor of tick microbial communities followed by average snowfall and 5-year average temperature (Fig. 2a). Temperature averaged over the previous year or 5 years were stronger predictors of tick microbiomes compared to the average temperature on the day of sampling. However, there was substantial variability across microbes (Fig. 2b). For example, NDVI was the most important variable for predicting the occurrence of *Ralstonia*, whereas snowfall and temperature were more important for *Massilia* and *Bacteroides* ASVs (Fig. 2b). Host sex was not as important across the microbial communities compared to microclimatic variables (Fig. 2a), but, for example, was the most important predictor for *Rickettsia* (Fi. 2b). ALE plots of the conditional relationship of each predictor support the importance of non-linear microbial responses and inter-taxa variability (Fig. 2 c & d). The ASVs that most strongly responded to the average temperature gradient and maximum snow depth showed either strong non-linear decreases (medium dashed lines, non-linear increases (solid lines) or a ‘u-shape’ with declining and increasing (small dashed lines) in occurrence probabilities (Fig. 2c & d). For example, the average probability of sampling *Francisella* steeply increased from ~0.2 to 0.5 °C when yearly average day temperature was > 13 °C (Fig. 2d, holding all other variables constant). However, the probability of sampling *Ralstonia* decreased from 0.6 to 0.25 when average temperature exceeded < 5 °C (Fig. 2c). Generally, our analysis pointed to an average yearly temperature of~5 °C and ~13 °C and ~250 and ~375 mm maximum snowfall as putative thresholds for changes in the tick microbiome (Fig. 2 c/d). We caution that due to the highly correlated nature of these microclimate predictors that these relationships could be proxies for the mechanistic drivers of microbiome change in *I. scapularis*.

We further examined the predictors that shaped *Borreliella* occurrence and found that the 5-year average temperature followed by vegetation cover (NDVI) had the highest importance scores (Fig. 3a). ICE plots revealed that higher temperature increased the probability of *Borreliella* at > 6 °C yet this effect plateaued rapidly (Fig. 3b). In contrast, NDVI values > 70 decreased the probability of observing *Borreliella* in these ticks (unlike what we found at the community level, Fig. S4). In each case, these variables had much less impact on ticks with a high predicted baseline probability of *Borreliella* occurrence (i.e., probabilities > 0.9, Fig. 3b & c).

**Fig. 3:**
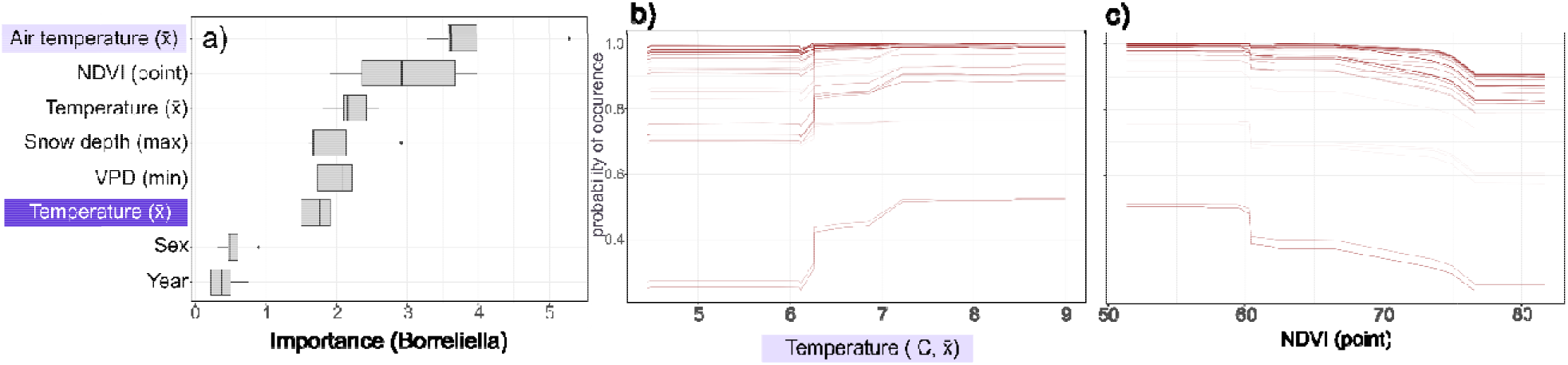
**a)** Variable importance and b/c) Individual conditional expectation (ICE) plots for the a) fiver year average temperature and b) NDVI (vegetative cover). Individual lines reflect change in microbe occurrence probabilities within individual ticks while holding all other variables constant in the model.

### Including microbial associations

Our models just including microbial associations could increase the predictive power by on average 70% compared to our MrIML approach, with our MRF model able to correctly predict the presence of at ASV on average ~85% of the time (Fig. 4a). The association only MRF model also had a similar performance compared to our spatial CRF model which included the environmental and host predictors. Adding spatial relationships decreased the specificity of the MRF model (Fig. 4b) but improved the sensitivity of the model (Fig. 4c). Adding spatial relationships did not substantially alter model predictive performance of the CRF models (Fig. 4). The CRF models had marginally better performance for predicting the absence of ASVs with an average sensitivity of ~0.45 compared to ~0.40 for the non-spatial MRF model (Fig. 4c).

**Fig. 4:**
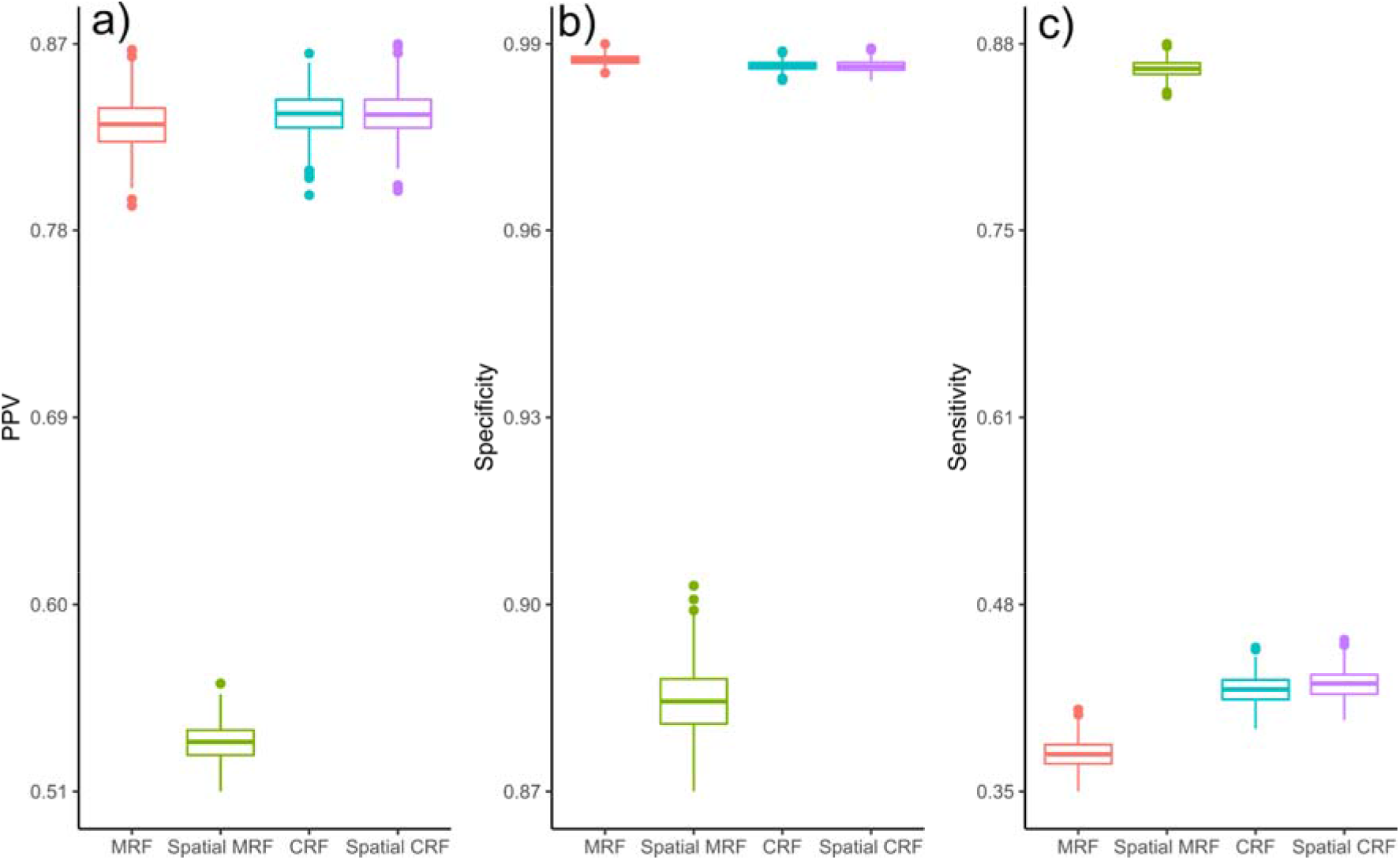
Model performance across graphical network models. A) Positive predictive value (PPV), b) specificity and c) sensitivity across MRF and CRF models.

### Tick microbiome co-occurrence network

When we further interrogated the spatial CRF co-occurrence network, we found that strong positive microbial associations (coefficients > 0.1) between ASVs were by far more common compared to negative associations (Fig. 5). We chose to focus on the spatial CRF model as this model limits ‘false’ associations between ASVs that were a product of shared environmental and or spatial proximity (N. Clark pers. Com). To simplify the network, we filtered edges with positive coefficients ≥ 0.4 and identified several taxa that may play a more important role in tick microbiome composition (i.e., microbiome hubs). *Bacteroides* had the highest degree (12 coefficients > 0.4) followed by *Corynebacerterium.1*, (11 coefficients > 0.4, Table S2). The positive association network also revealed one main connected cluster as well as disconnected pairs of ASVs (Fig. 5a). The main cluster included *Cornybacterium.1*, *Lawsonella*, *Sphingomonas*, *Pseudomonas*, and *Mycobacterium*. One disconnected pair included *Streptococcus* and *Borreliella* and further examination of the coefficients showed this relationship was bidirectional (i.e., presence of *Streptococcus* increased the probability of *Borreliella* occurrence and *vice versa*, Table 1). Our focal taxa were also represented in the negative association network with, for example, a strong negative association (CRF coefficient = −0.28) between *Ehrlichia* and *Mycobacterium* (Fig. 5b, Table 1). Interestingly, this relationship appears to be unidirectional as the occurrence of *Mycobacterium* was not associated with *Ehrlichia* (Table 1). We also detected negative associations involving *Francisella* and *Borrelia* (Fig. 5b), but further examination of the coefficients revealed that these negative associations did not impact the probability of occurrence of these pathogens (Table 1).

**Fig. 5:**
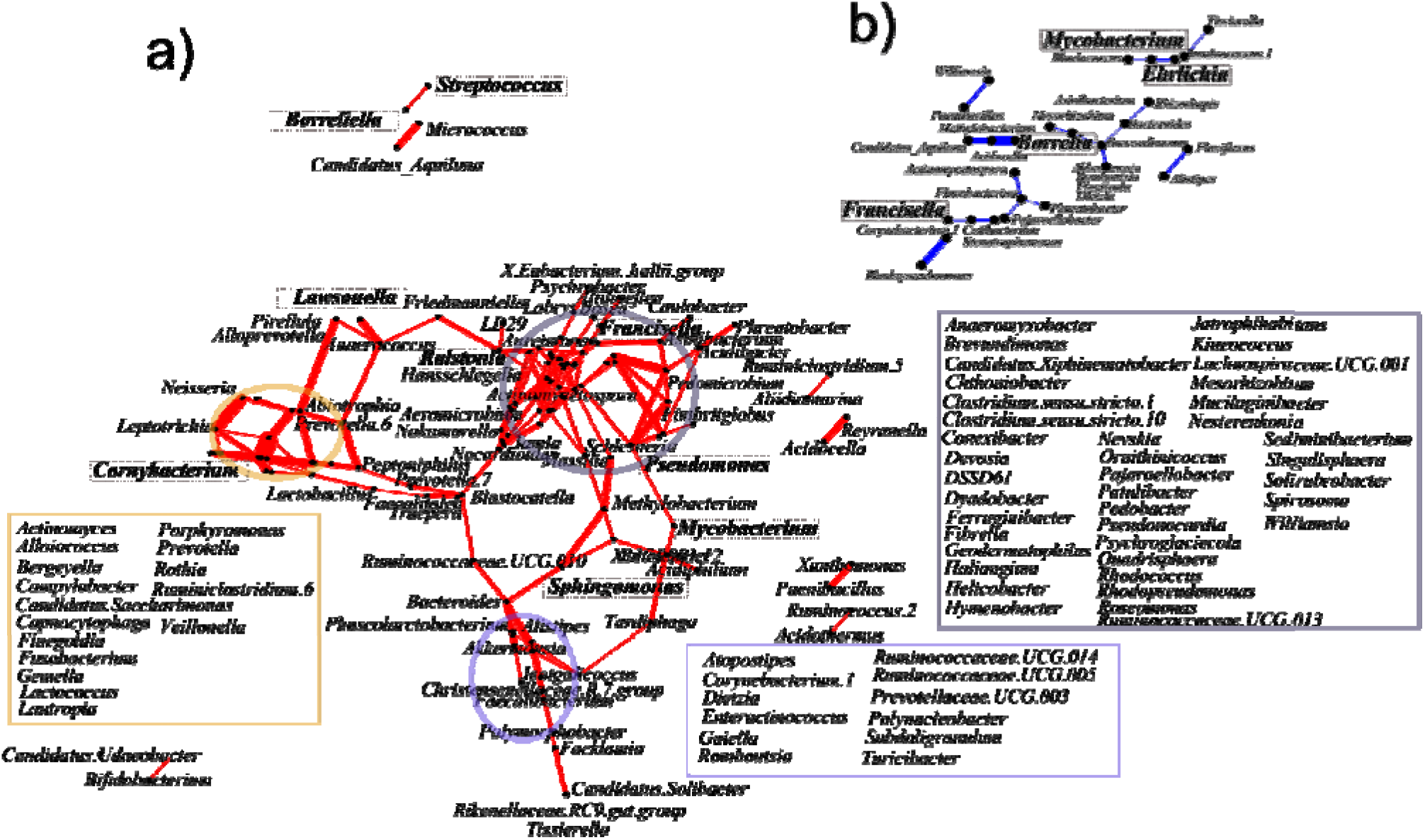
Positive (red, a) and negative (blue, b) association networks based on the spatial CRF model. Nodes (vertices) are positioned using multidimensional scaling (MDS) and represent amplicon sequence variants (ASVs). Nodes closer together have a more similar association profile. Edge widths are scaled by the strength of the association. To aid interpretability, coloured boxes show the taxa within each corresponding oval. ASVs in bold are the focal taxa. a) Positive associations where the coefficient > 0.4. b) Negative associations where the coefficient >-0.1. ASVs are in italics. ASVs with coefficients below these threshold values were removed. The network was qualitatively similar for the spatial MRF model.

### Individual indicator taxa

For the taxa of interest, combining MrIML and CRF results further highlighted the importance of microbial associations in predicting tick microbiome dynamics. Environment alone could explain the occurrence of 8/14 focal taxa reasonably well (MrIML PPV < 0.5, Table 1). We could, for example, predict the occurrence of *Mycobacterium* and *Borreliella* correctly 100% and 95% of the time respectively using environmental/host predictors (PPV = 1.0/0.95, Table 1). However, in the CRF model, environment and host variables were not the most important indicating that non-linear relationships are important (MRF/CRF models use logistic regression to estimate coefficients). The MrIML and CRF models did in some cases provide similar inference. For example, both approaches confirmed tick sex as the most important predictor of *Rickettsia* (Fig. 2b, Table 1). Both modelling approaches also identified that vegetation, as measured by NDVI, had a notable effect on *Borreliella* occurrence with a negative association with increasing NDVI (Table 1, Fig. 3c). For the taxa poorly predicted by environment, our spatial CRF model identified that associations with other ASVs were more important (high CRF importance and coefficients >0.1, Table 1). For example, the occurrence of *Lawsonella* within ticks was poorly predicted by the environment alone (PPV = 0, or in other words our model could never correctly predict the occurrence of *Lawsonella*). However, in the CRF model, we found the presence of *Anaerococcus* was the dominant variable predicting the occurrence of *Lawsonella* (relative proportional importance = 0.96, CRF coefficient = 1.01). Our CRF model also identified the importance of site effects for *Francisella*, *Ralstonia*, and *Sphingomonas* not captured in the MrIML models. Moreover, our CRF model identified a few important interactions shaping the occurrence of microbes with, for example, the presence of *Neisseria* at higher temperatures being relatively important for the occurrence of *Corynebacterium.1* (Table 1).

## Discussion

Our study highlights the important role that positive microbial associations play in shaping microbiome composition within *I. scapularis*. Variables including temperature were important for some taxa, including *Borreliella*, however, overall these ecological processes played a much more minor role in shaping these communities at this scale. Even though they were not as common, our approach also quantified some important negative associations including between potentially pathogenic taxa. Further, this study provides a case study to how two complimentary machine learning-based approaches can help untangle how microbial associations, host, and environment can shape microbiomes. We stress that even though these findings are necessarily associative, they can provide new hypotheses on how pathogens and symbionts might interact within tick species and how these relationships can be shaped by the environment.

Our approach enabled us to quantify the relative dominance of positive microbial associations compared to other ecological processes in shaping microbiome assembly in *I. scapularis*. However, there were still some important effects of the environment on microbial occurrence. For example, we found that yearly and five-year average temperature was important for a variety of bacteria including symbionts and pathogens. Temperature is known to be particularly important for questing (i.e., actively seeking hosts) and subsequently for pathogen acquisition [40–42], and our study shows that it is important for the tick acquiring other potential symbiotic or commensal microbes as well. The relationship between temperature and the presence of *Borreliella* is particularly intriguing and may have implications for the observed range expansion of *I. scapularis* [43] and increase in *B. burgdorferi* prevalence over time [44] in ticks in the Upper Midwest. Laboratory studies have previously demonstrated that infection of *Ixodes* spp. ticks by *B. burgdorferi* can enhance tick survival and alter questing behavior based on temperature and humidity. Herrmann and Gern [45] found that infection with *B. burgdorferi s.l*. increased *I. ricinus* nymphal and adult survival under “challenging” temperature and humidity conditions, independent of tick behavior. However, we also showed that increasing temperature decreased occurrence probability for some putative non-pathogenic taxa (Fig. 2c). The nonlinear declines in occurrence for some taxa are possibly a result of shifts in soil microbiomes associated with taxon shifts in warming conditions. Even though we controlled for soil contamination, soil microbial communities play a role in tick microbiome composition [46].

The responses to maximum snowfall and NDVI were also highly divergent across ASVs. For example, higher NDVI values increased microbiome turnover overall, yet for some species, such a *Borrelielia* and *Corynebacterium. 1* the probability of occurrence decreased with NDVI. NDVI has a positive relationship with plant diversity [e.g., 47] and small mammal diversity [e.g., 48] and the ‘dilution’ of host species may underlie the negative relationship we detected. *B. burgdorferi .s.l* prevalence in European *I. ricinus* ticks has been previously found to decline with plant richness but increase with vegetative density [49]. While *I. scapularis* is largely limited to forested habitat in the Upper Midwest, areas of high NDVI may harbour increased mammal species richness and include more *Borreliella* refractory reservoirs, such as white-tailed deer (*Odocoileus virginianus*), Virginia opossums (*Didelphis virginiana*), or raccoons (*Procyon lotor*) [50, 51].

Overall microbial associations were, however, a much stronger predictor of *I. scapularis* microbiomes. While few studies have quantified the relative importance of these microbial associations, positive associations have found to be much more common relative to negative associations in European *I. ricinus* microbiomes [10, 11] and *Ixodes pacificus* in Southern California [52]. The dominance of positive associations may be due to the importance of facilitation as a process in these communities and may also be signature to parallel colonization of ticks (i.e., ticks get colonized by groups of microbes) [10]. *Corynebacterium.1* and *Bacteroides* were putative hubs potentially playing an important role in the tick microbiome. *Bacteroides*, for example, is known to be a crucial taxa in the human microbiome (a ‘keystone species’) also having many associations with other microbes [e.g., 53], but why it plays an important role in the tick microbiome warrants further investigation.

While the pathogens were less connected in the network, we did detect positive and negative interactions between potentially pathogenic and non-pathogenic taxa. For example, we identified that a *Fusobacterium* was by far the most important predictor *of Anaplasma* occurrence (Table 1) and this link is supported by other studies. For example, in the spleen of *Anaplasma*-positive wild small mammals, *Fusobacterium* was found to be upregulated [54]. The link between *Anaplasma* and *Actinomyces*, however, is unknown currently. In contrast, *Borreliella* was much less connected in our network compared to what has been found in *I. ricinius* [10, 55]. Some studies have shown that *B. burgdorferi* occurrence had little impact on patterns of tick microbiome composition and diversity [e.g., 56], while other have shown disruptive effects of *B. burgdorferi* infection [11] This discrepancy could be due to differences in *Ixodes* or *Borreliella* species observed in the referenced studies as associations between microbes can be even genotype specific [10, 57]. The strongest association noted with *Borreliella* in this study was with *Streptococcus*. *Streptococcus* has been rarely found in previous *I. scapularis* microbiome studies but has been noted in wild nymphs [58] and lab-reared larvae [18]. In our samples it was more abundant, with 23 ticks (13 females, 10 males) producing more reads than the controls. 9 of these originated from a single collection site (Camp Ripley). Further observational and manipulative experiments on both adult and nymphal ticks are needed to resolve the relationship of *B. burgdorferi s.l*. species to the microbiome of their host ticks.

While including the geographic distance between sites in our model alone led to a decrease or had no effect in mean PPV in our MRF and CRF models respectively, it did increase sensitivity (i.e., the model could better distinguish where the ASVs occurred Fig. 3). Spatial effects have been found to be important in mammal microbiomes [7, 8, 59], and our findings suggest microbial dispersal may also play a role in tick microbiomes, yet our results were not conclusive. Ticks and their microbes can only disperse smaller distances [60] and including finer spatial sampling within sites may help untangle the relative role of dispersal acting on the composition of tick microbiomes. For some taxa, including *Francisella*, site was the only significant factor and these taxa were independent of other microbes. *Francisella* may be pathogenic but non-pathogenic *Francisella*-like endosymbionts are known from a variety of tick genera including *Amblyomma*, *Dermacentor*, and *Ornithodoros*. *Francisella* is not normally associated with *Ixodes*. Further identification of this bacterium to determine pathogenicity was beyond the scope of this study.

While our study provides insight into the drivers of *I. scapularis* microbiome dynamics, our findings are necessarily associative and should help guide future more mechanistic studies [36]. These studies could include the controlled examination of host seeking behaviour under varying environmental conditions using ticks that have been infected with pathogens of interest. While some experiments have looked at behavioural changes related to infection of *I. scapularis* by *B. burgdorferi* [61], the associations between the microbiome and tick behaviour warrants further examination. Similarly, the microbiome may affect tick survival under adverse environmental conditions. Laboratory studies have shown that infection of ticks by *A. phagocytophilum* enhances tolerance to freezing by stimulating antifreeze glycoprotein production [62]. Although our model found no evidence relating *A. phagocytophilum* prevalence to site temperature, this association, and other, unknown microbiome-tick interactions, could influence tick landscape distribution.

Our ability to interpret the positive and negative associations between taxa we detected could be further improved by extracting samples from specific tick organs. For, example, quantifying organ-specific microbial communities can help untangle if putative associations are direct (e.g., microbes compete directly for resources) or indirect (i.e., the presence of a microbe indirectly impacts colonisation of other microbes via immune response [63]). While we rigorously washed ticks and quantified soil microbes to help focus our analysis on tick-specific microbes, our sampling was unable to exclude organisms present in the tick’s previous blood meals. Further, we did not quantify differences in blood meals across ticks that can shape the diversity and composition of tick microbiomes [64]. However, as we excluded rare ASVs, and the number of potential host species in the Upper Midwest US is limited, differences in blood meal are less likely to impact our results. Additionally, examination of the microbiome community in ticks fed on different host animals did not appear to alter community composition [65]. Lastly, while our MrIML and CRF models provide complementary insights into community dynamics, directly comparing predictive performance across models is not possible as uncertainty is not quantifiable and site effects are not captured in MrIML models. Future versions of MrIML will have functionality to calculate uncertainty in predictive performance across training and testing data splits as well as to calculate associations between response variables [27].

We show that associations between microbes play an important role in shaping microbiome dynamics in *I. scapularis* in the Upper Midwest, yet the importance of this ecological driver is likely to extend across continents and tick species. The distribution of important pathogenic groups such as *Anaplasma* were associated with the presence of putatively symbiotic groups within ticks. Other genera were more correlated with differences in climate. Studies like ours that can start to untangle these complex relationships promise to guide mechanistic experiments and better predict the distribution of tick pathogens and symbionts alike into the future.

## Acknowledgements

This project was supported by a grant from the MN Futures program of the University of Minnesota. NFJ was supported by an Australian Research Council Discovery Project Grant (DP190102020).

