## Supplementary material for "Positive associations matter: microbial relationships drive tick microbiome composition": Fig S1-4

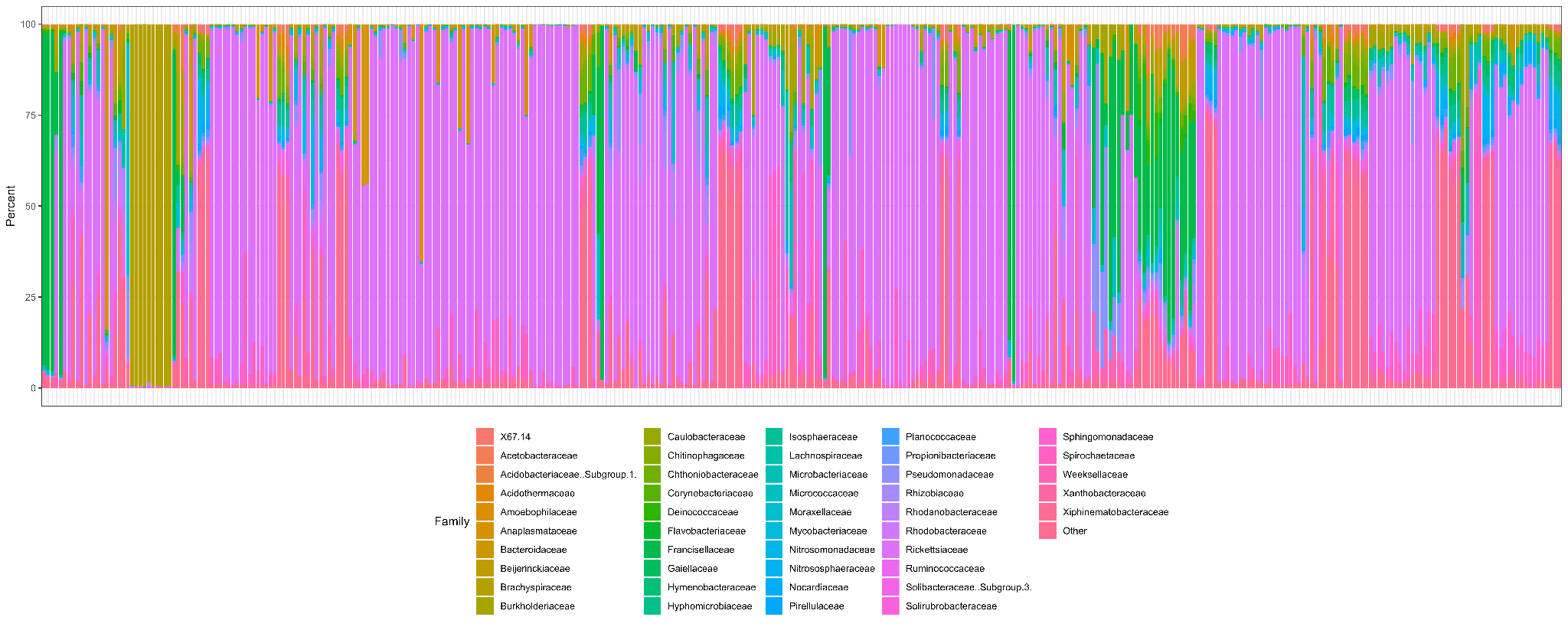

**Fig. S1:** Family-level taxonomic assignment across the 335 ticks sampled in this study. Note that columns that are almost entirely light brown on the left are the spatter control wells that comprised almost entirely of Brachyspiraeae.

**Table S1:** Mr IML model performance across random forest (RF), logistic regression (GLM) and extreme gradient boosting (XGB) algorithms.

| **Model** | **Average PPV** | **Average specificity** | **Average sensitivity** |
| --- | --- | --- | --- |
| **Random Forests (RF)** | **0.15** | **0.89** | **0.95** |
| Logistic regression (GLM) | 0.09 | 0.84 | 0.94 |
| Extreme gradient boosting (XGB) | 0.07 | 0.84 | 0.94 |

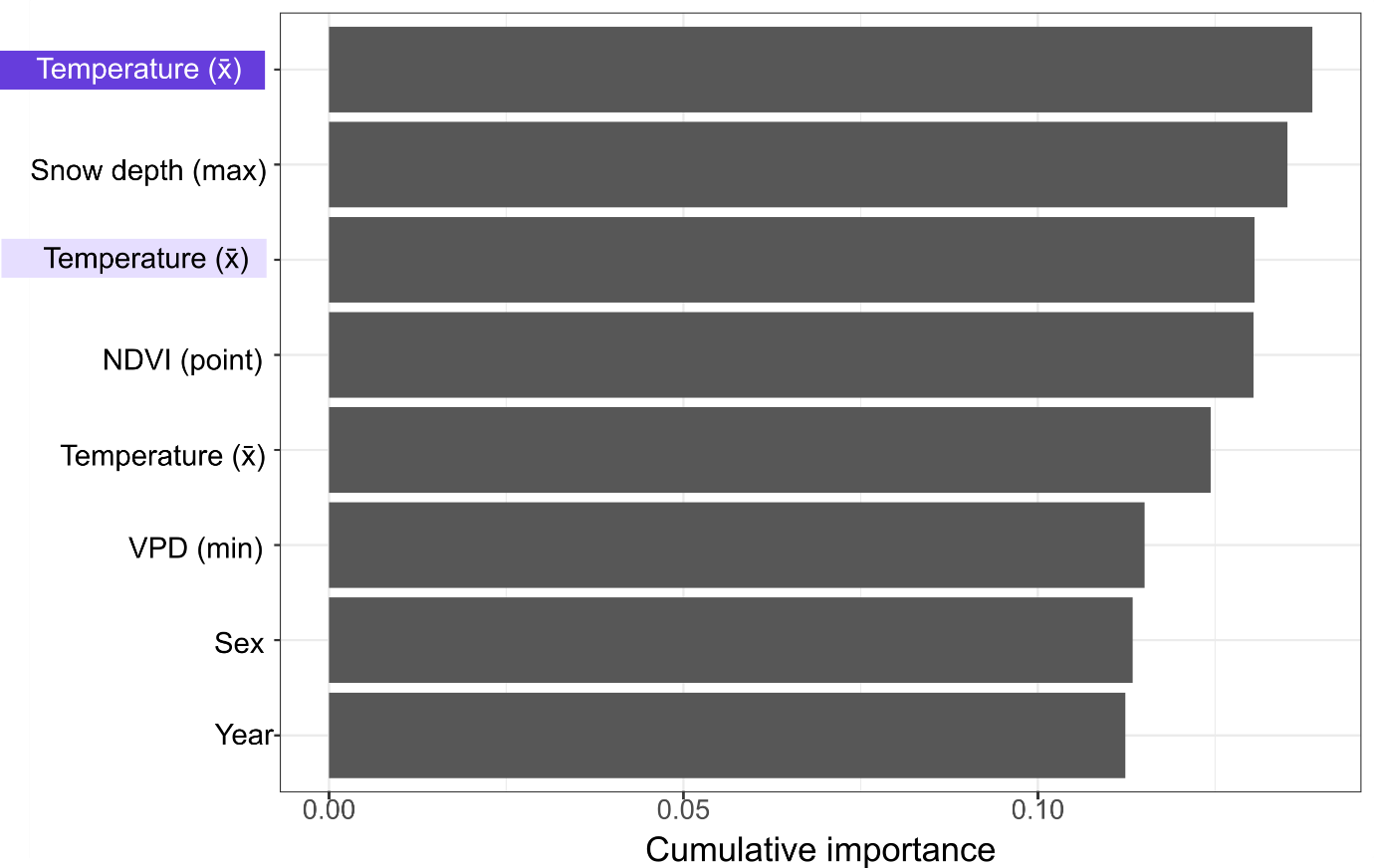

**Fig. S2:** Variable importance plot for our GLM-based linear model.

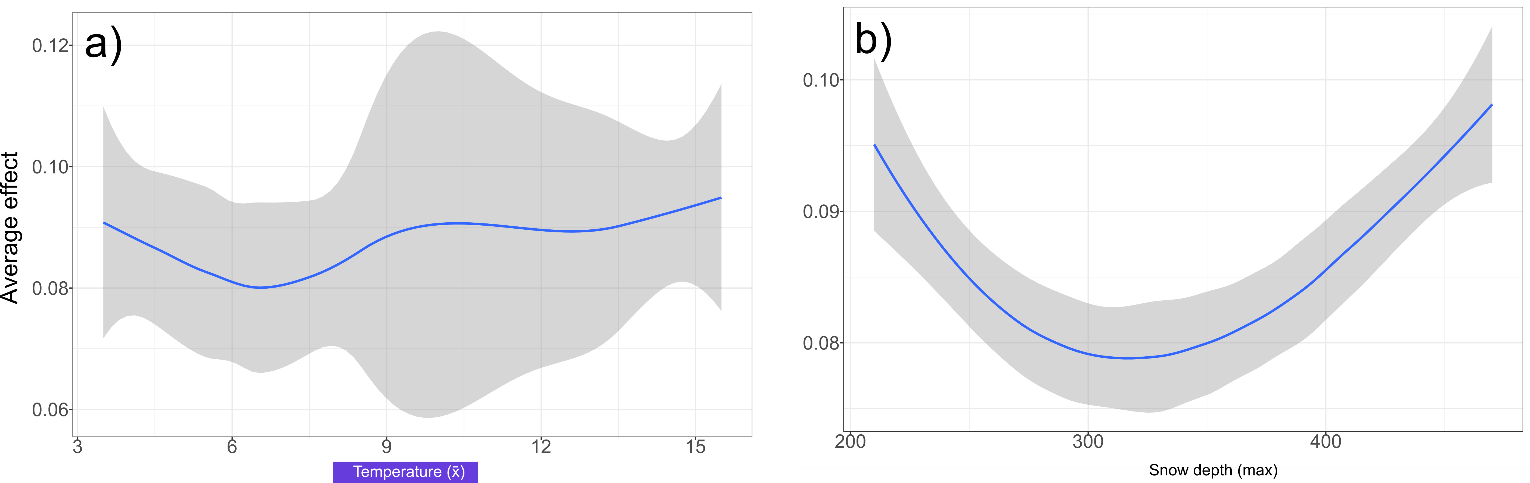
 **Fig. S3:** Global accumulated local effects plots showing the average relationship between a) mean yearly temperature and b) maximum snow fall for 231 ASVs commonly found in our tick samples.

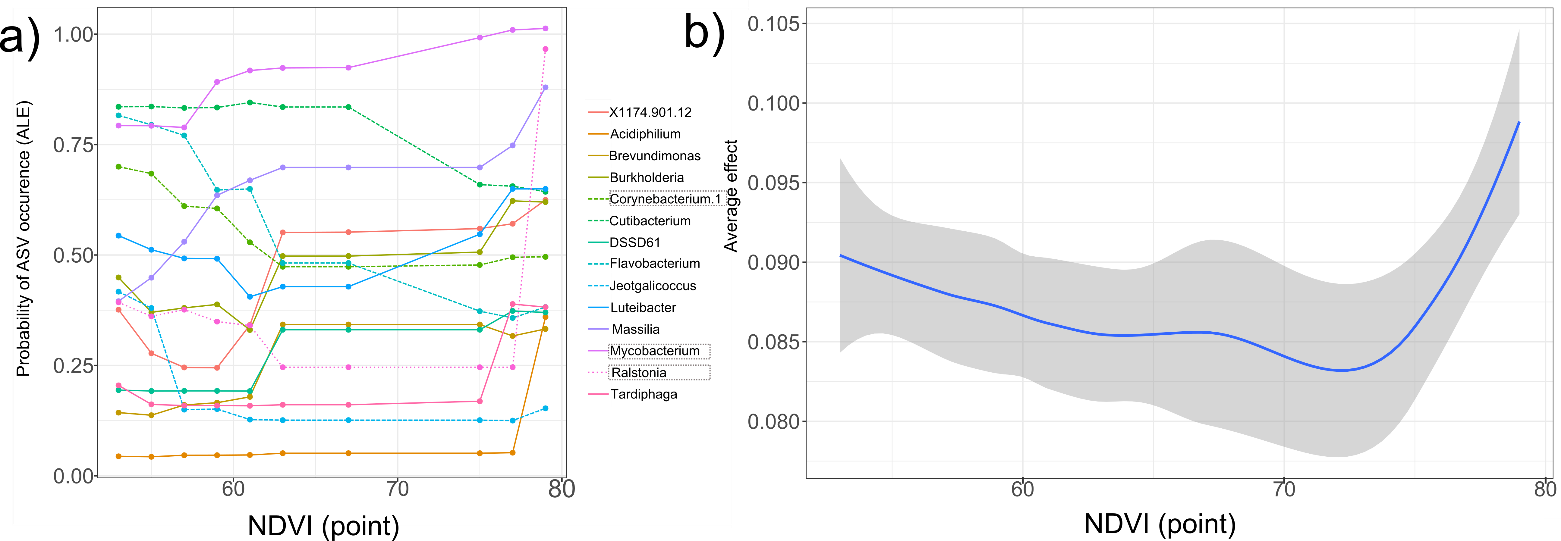

**Fig. S4:** Accumulated local effects plots for the a) the individual ASVs and b) community-wide (‘global’) responses to the normalized difference vegetation index (NDVI) across our sites. Focal taxa are indicated by the dashed boxes. Medium dashed lines =ASVs with a generally negative relationship with NDVI. Small-dashed lines = decline followed by increase. Unbroken = positive relationship with NDVI.

.

**Table S2:** Positive association (coefficient >0.4) degree for each taxa (ASVs with degree=0 not included). D = degree. Bold text are the focal taxa.

| **ASV** | **D** | **ASV** | **D** | **ASV** | **D** | **ASV** | **D** | **ASV** | **D** | **DASV** | **D** |
| --- | --- | --- | --- | --- | --- | --- | --- | --- | --- | --- | --- |
| Bacteroides | 12 | X1174.901.12 | 4 | Jeotgalicoccus | 3 | Lactococcus | 2 | Anaeromyxobacter | 1 | Phascolarctobacterium | 1 |
| Corynebacterium.1 | 11 | Abiotrophia | 4 | Luteibacter | 3 | Lautropia | 2 | Bifidobacterium | 1 | Phreatobacter | 1 |
| Neisseria | 9 | Aeromicrobium | 4 | Massilia | 3 | Nakamurella | 2 | Borreliella | 1 | Pirellula | 1 |
| Hymenobacter | 8 | Capnocytophaga | 4 | Mesorhizobium | 3 | Nevskia | 2 | Bosea | 1 | Polymorphobacter | 1 |
| Solirubrobacter | 8 | Facklamia | 4 | Porphyromonas | 3 | Patulibacter | 2 | Candidatus.Saccharimonas | 1 | Polynucleobacter | 1 |
| Spirosoma | 8 | Jatrophihabitans | 4 | Prevotella.6 | 3 | Prevotella.7 | 2 | Candidatus.Solibacter | 1 | Prevotella | 1 |
| Pedomicrobium | 7 | Kineococcus | 4 | Rhodococcus | 3 | Prevotellaceae.UCG.003 | 2 | Candidatus.Udaeobacter | 1 | Psychrobacter | 1 |
| Bergeyella | 6 | Leptotrichia | 4 | Sphingomonas | 3 | Pseudomonas | 2 | Candidatus_Aquiluna | 1 | Psychroglaciecola | 1 |
| Blastocatella | 6 | Methylobacterium | 4 | Tardiphaga | 3 | Pseudonocardia | 2 | Caulobacter | 1 | Reyranella | 1 |
| Brevundimonas | 6 | Mycobacterium | 4 | Alistipes | 2 | Quadrisphaera | 2 | Christensenellaceae.R.7.group | 1 | Rikenellaceae.RC9.gut.group | 1 |
| Ferruginibacter | 6 | Roseomonas | 4 | Alloiococcus | 2 | Ralstonia | 2 | Clostridium.sensu.stricto.1 | 1 | Ruminiclostridium.5 | 1 |
| Geodermatophilus | 6 | Sediminibacterium | 4 | Alloprevotella | 2 | Rhodopseudomonas | 2 | Clostridium.sensu.stricto.10 | 1 | Ruminiclostridium.6 | 1 |
| Mucilaginibacter | 6 | Actinomyces | 3 | Atopostipes | 2 | Rothia | 2 | Enteractinococcus | 1 | Ruminococcaceae.UCG.005 | 1 |
| Nocardioides | 6 | Akkermansia | 3 | Candidatus.Xiphinematobacter | 2 | Ruminococcaceae.UCG.010 | 2 | Fibrella | 1 | Ruminococcaceae.UCG.013 | 1 |
| Pedobacter | 6 | Asinibacterium | 3 | Corynebacterium | 2 | Singulisphaera | 2 | Francisella | 1 | Ruminococcaceae.UCG.014 | 1 |
| Schlesneria | 6 | Aureimonas | 3 | Dietzia | 2 | Williamsia | 2 | Gaiella | 1 | Ruminococcus.2 | 1 |
| Anaerococcus | 5 | Burkholderia.Caballeronia.Paraburkholderia | 3 | Faecalibacterium | 2 | X.Eubacterium..hallii.group | 1 | Iamia | 1 | Streptococcus | 1 |
| Chthoniobacter | 5 | Campylobacter | 3 | Faecalitalea | 2 | Acidibacter | 1 | Lachnospiraceae.UCG.001 | 1 | Subdoligranulum | 1 |
| DSSD61 | 5 | Conexibacter | 3 | Friedmanniella | 2 | Acidiphilium | 1 | Lawsonella | 1 | Tissierella | 1 |
| Dyadobacter | 5 | Devosia | 3 | Gemella | 2 | Acidocella | 1 | LD29 | 1 | Truepera | 1 |
| Pajaroellobacter | 5 | Fimbriiglobus | 3 | Hansschlegelia | 2 | Acidothermus | 1 | Micrococcus | 1 | Turicibacter | 1 |
| Peptoniphilus | 5 | Finegoldia | 3 | Helicobacter | 2 | Actinomycetospora | 1 | Nesterenkonia | 1 | Xanthomonas | 1 |
| Romboutsia | 5 | Fusobacterium | 3 | Labrys | 2 | Aliidiomarina | 1 | Ornithinicoccus | 1 |  |  |
| Veillonella | 5 | Haliangium | 3 | Lactobacillus | 2 | Aliihoeflea | 1 | Paenibacillus | 1 |  |  |
